# Enhanced germination and electrotactic behaviour of *Phytophthora palmivora* zoospores in weak electric fields

**DOI:** 10.1101/2023.01.19.524801

**Authors:** Eleonora Moratto, Stephen Rothery, Tolga O. Bozkurt, Giovanni Sena

## Abstract

Soil-dwelling microorganisms use a variety of chemical and physical signals to navigate their environment. Plant roots produce endogenous electric fields which result in characteristic current profiles. Such electrical signatures are hypothesised to be used by pathogens and symbionts to track and colonise plant roots.

The oomycete pathogen *Phytophthora palmivora* generates motile zoospores which swim towards the positive pole when exposed to an external electric field *in vitro*.

Here, we provide a quantitative characterization of their electrotactic behaviour in 3D. We found that a weak electric field (0.7 - 1.0 V/cm) is sufficient to induce an accumulation of zoospore at the positive pole, without affecting their encystment rate. We also show that the same external electric field increases the zoospore germination rate and orients the germ tube’s growth. We conclude that several early stages of the *P. palmivora* infection cycle are affected by external electric fields.

Taken together, our results are compatible with the hypothesis that pathogens use plant endogenous electric fields for host targeting.

## Introduction

Soil-dwelling organisms such as bacteria, fungi and plant roots use environmental cues of physical, chemical and biological origin to navigate through this complex environment. Plant pathogens and symbionts must identify their host roots in soil to initiate biological interactions. They achieve this through the detection of chemical and electrical cues produced by plant roots (Steinkellner et al., 2007). While the role of chemical signalling is well understood, little is known about the role of electrical cues in this context.

Electrotaxis, or galvanotaxis, is the change of an organism’s direction of migration in response to an electric field (Dineur, 1892). Electrotropism, or galvanotropism, is the change of an organism’s direction of growth in response to an electric field (Gilroy, 2008; Muthert et al., 2020). Electrotaxis and electrotropism have been documented in animals (Clarke et al., 2013; Crampton, 2019), plants (Muthert et al., 2020; Salvalaio et al., 2022), fungi (McGillivry & Gow, 1986) and oomycetes (Miller & Gow, 1989). Early studies have shown that zoospores of several plant pathogens respond to weak electric fields *in vitro* by swimming preferentially towards one of the poles (Khew & Zentmyer, 1974).

Roots of several plant species are known to generate electric fields (Toko et al., 1989) with current densities between 3 and 27 mA/m^2^ (Miller et al., 1991; Miller & Gow, 1989; Ryan et al., 1990; Taylor & Bloom, 1998). This suggests that some pathogen zoospores could use the root’s endogenous electric signature as a target (van West et al., 2002).

The oomycete pathogen *Phytophthora palmivora*’s infection cycle begins with swimming zoospores that identify the host root (Figure 1A). This is followed by encystment, which includes the loss of zoospore flagella and attachment to the host (Judelson & Blanco, 2005). The last stage of the early infection cycle is zoospore germination (Figure 1A) which is then followed by plant tissue invasion.

**Figure 1:**
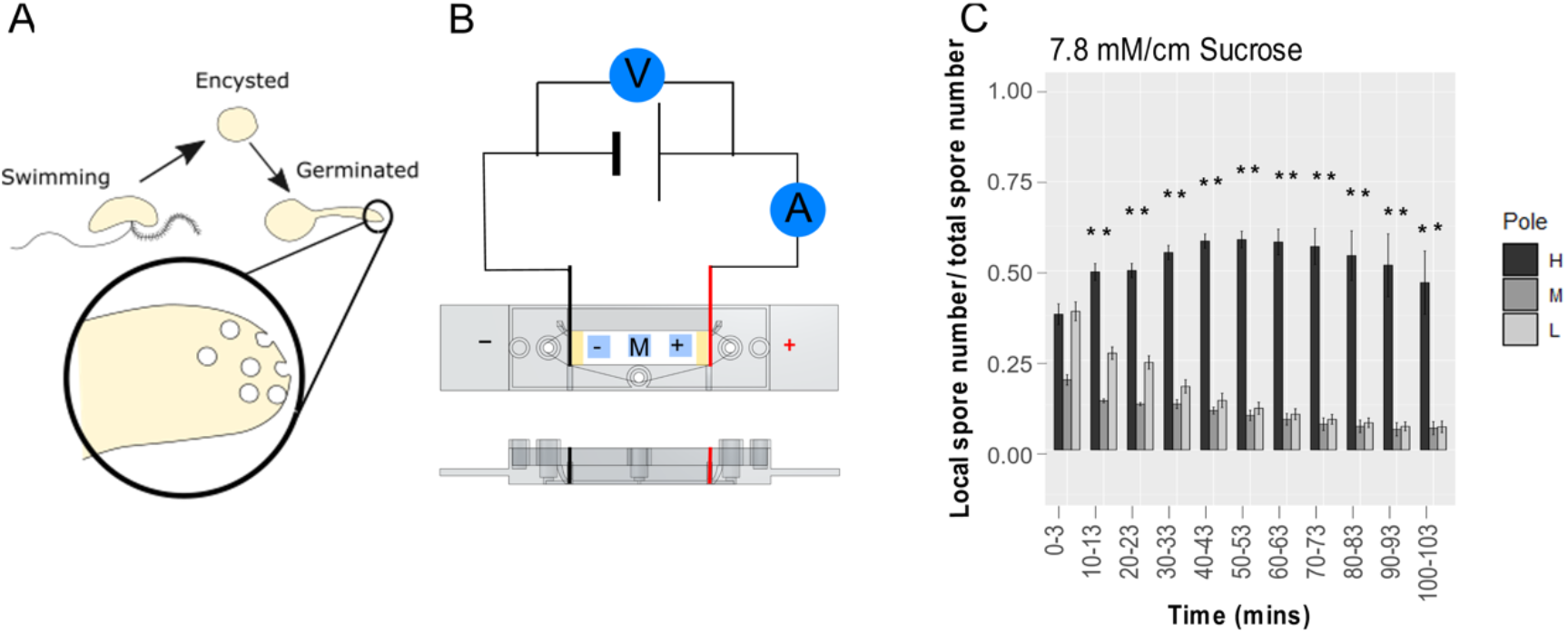
Assay for zoospore electrotaxis. **A)** Diagram of early *P. palmivora* zoospore life cycle stages.Swimming *P. palmivora* spores are characterised by two flagella used for propulsion; encysted spores lose their flagella and undergo organelle rearrangement, causing a change of shape from bean-like to almost spherical; germination is characterised by the emergence of the germ tube; vesicles fuse with the plasma membrane at the tip of the germ tube allowing for directional growth **B**) Schematic of the 3D-printed V-slide set-up for *P. palmivora* electrotaxis experiments. The black and red electrodes are connected to an external power supply; the three imaging regions are marked −, M, +; V, voltmeter; A, ammeter. **C)** Time-course of the distribution of spores exposed to a strong sucrose gradient. At each time-point, each bar represents the proportion of swimming spores in the slide’s regions with high (H, dark grey), middle (M, grey) and low (L, light grey) sucrose concentration; error bars, s.e.p. After only 10 minutes, the proportion of spores in the high-sucrose concentration region becomes significantly higher than the one in the low-concentration region (** = p-value < 0.01 between H and L; t-test, n = 5).

*P. palmivora* zoospores are electrotactic towards the positive pole *in vitro* (Morris et al., 1992). However, we lack a full understanding of the dynamics of zoospore responses in a 3D environment when exposed to an electric field. Moreover, although electric fields are known to change germination and hyphal orientation in some mycelial organisms (McGillivry & Gow, 1986), the effect of electric fields on the various stages of the *P. palmivora* life cycle remains largely unknown.

In this paper, we describe a novel setup to monitor *in vitro* the dynamics of a population of *P. palmivora* zoospore when exposed to a constant electric field. We characterize their movement in 3D and their transition to encystment, (Judelson & Blanco, 2005), and germination (Figure 1A).

## Results

### Developing a quantitative assay to characterize zoospore electrotaxis in 3D

To study zoospore electrotaxis in three dimensions, we developed a setup to expose the spores to a constant electric field and monitor their position over time. Briefly, we pipetted the spores in the middle of an *ad-hoc* 3D-printed chamber (V-slide, Figure 1B and Supplementary Figure 1) fit with a standard glass coverslip at the bottom, two platinum foils as electrodes and filled with buffered liquid medium. To minimize the formation of chemical gradients through electrochemical reactions, the electrodes were coated in agarose gel. An automated inverted fluorescence microscope was used to capture a time series of Z-stack images in the middle of the slide and close to each electrode (see Methods for details). A macro was written in ImageJ to quantify spore numbers in each image.

To test the setup and validate the image processing method, we first exposed the zoospores to a chemical attractant gradient and quantified the change in zoospore distribution across the V-slide. To this goal, we added a small amount of sucrose, a known zoospore attractant (C. Bimpong et al., 1969), to one side of the chamber to generate a concentration gradient of 7.8 mM/cm. As expected, after only 10 minutes we observed an increase in zoospore concentration on the side of highest sucrose concentration (Figure 1C; t-test, p<0.01). To quantify the accuracy of our automated routine for spore-counting, we counted by hand the true spores which were misidentified by the macro (i.e. false negatives). Overall, swimming spores were misidentified less than 2% of the time while encysted spores were more likely to be missed by the macro (7.73% false negatives) (Supplementary Table 1).

### Exposure to an external electric field changes zoospore swimming dynamics in three dimensions

We pipetted the zoospores in the top layer of the middle region of the V-slide and analysed their response to constant electric fields with intensities 0.5, 0.7 and 1.0 V/cm (Figure 2). At any given time and in all tested field intensities, most of the swimming spores were detected in the top layer of the liquid media, while some remained in the central volume and others precipitated at the bottom when they lost their flagella (encystment) (Supplementary Figure 2).

**Figure 2:**
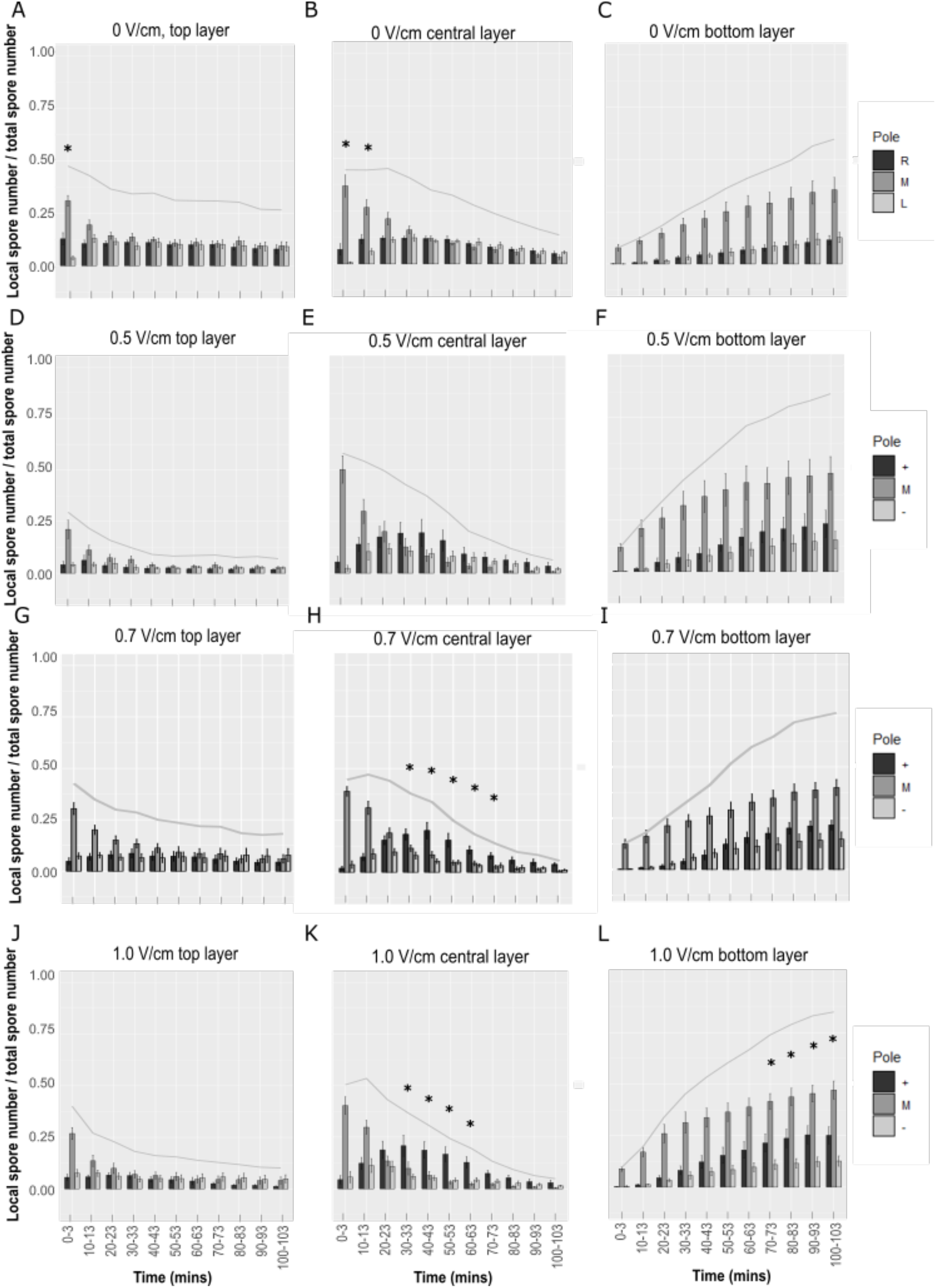
Zoospores are electrotropic towards the positive electrode. Time-course of the distribution of spores detected in the top, central and bottom layers of the medium in the V-slide. At each timepoint, each bar represents the proportion of spores close to the positive electrode or the right side (dark grey), the negative electrode or the left side (light grey), and in the middle (medium grey); error bars, s.e.p. The grey line is the proportion of all spores detected in that layer, at each timepoint. The spores in the central layer accumulated at the positive pole when exposed to 0.7 and 1.0 V/cm approximately 30 to 70 minutes after the start of the field (H & K; ^*^= p-value < 0.05 between positive and negative regions; t-test, n = 9).

To maintain sufficient spatial resolution, we organised the analysis in three vertical layers (top, central and bottom) and three horizontal regions (Right or Positive pole, Middle, Left or Negative pole). We identified the top layer as the first 540μm of the liquid media (Methods and Supplementary Figure 2), the bottom layer as the single slice where encysted spores are in focus and the central layer as all the volume in-between.

Without an electric field, the swimming spores in the top and middle layer quickly diffused symmetrically from the initial middle region to reach a uniform distribution (Figure 2 A & B and Supplementary Figure 3 A). Encysted spores gradually accumulate on the bottom layer while maintaining the initial bias on the middle region where they had been pipetted at the start of the experiment (Figure 2 C and Supplementary Figure 3 A).

When exposed to 0.5 V/cm, the apparent accumulation of spores to the positive pole is not statistically significant (Figure 2 D & E and Supplementary Figure 3 B). With 0.7 V/cm, after 40 to 70 minutes we observed in the middle layer a statistically significant accumulation of spores at the positive pole (Figure 2 H; t-test, p<0.05), but not in the top or bottom layers (Figure 2 G & I). At 1.0 V/cm, we observed similar behaviour in the top (Figure 2 J) and middle layer (Figure 2 K; t-test, p<0.05), while in the bottom layer the distribution was biased towards the positive pole (Figure 2 L; t-test, p<0.05).

We wondered if the observed accumulation at the positive pole was due to true electrotaxis or, instead, to a chemotactic response to a putative chemical gradient generated by electrochemical reactions in the medium. To address this, we pre-exposed the medium in the V-slide to 1.0 V/cm for 100 minutes, switched off the field, immediately pipetted the spores and imaged while maintaining the electric field off. We reasoned that if 1.0 V/cm was sufficient to generate a chemical gradient strong enough to be sensed by the spores, we would have observed chemotaxis in the V-slide. Instead, we found no difference in spore concentration between the positive and negative poles (Supplementary Figure 4), indicating that chemotaxis is not contributing in a detectable way to the spore dynamics observed in presence of electric fields, and that electrotaxis is the dominant mechanism.

### Ionic currents are required for zoospore electrotaxis

Early reports suggested that a pure electrostatic force could be sufficient to reorient the zoospore’s flagella, which might be charged (Morris & Gow, 1993). If this were the dominant mechanism of zoospore electrotaxis, we would expect the zoospore swimming dynamics to be independent of the ionic currents generated in the medium.

In the electrotaxis assay described above, a 1.0 V/cm field applied to our standard 1/500X MS medium generated an average (± s.e.m.) electric current of 56.6 μA ± 1.4 μA. To test the hypothesis above that zoospore electrotaxis depends on the field rather than the current, we repeated the same assay at 1.0 V/cm but now in deionised water, which resulted in a significantly lower 12.2 μA ± 0.5 μA current. Interestingly, no significant electrotactic behaviour was observed in these conditions (Figure 3), suggesting that the ionic currents in the medium, and not just the electric field applied, play an important role in the biophysical mechanism of electroreception in zoospores.

**Figure 3:**
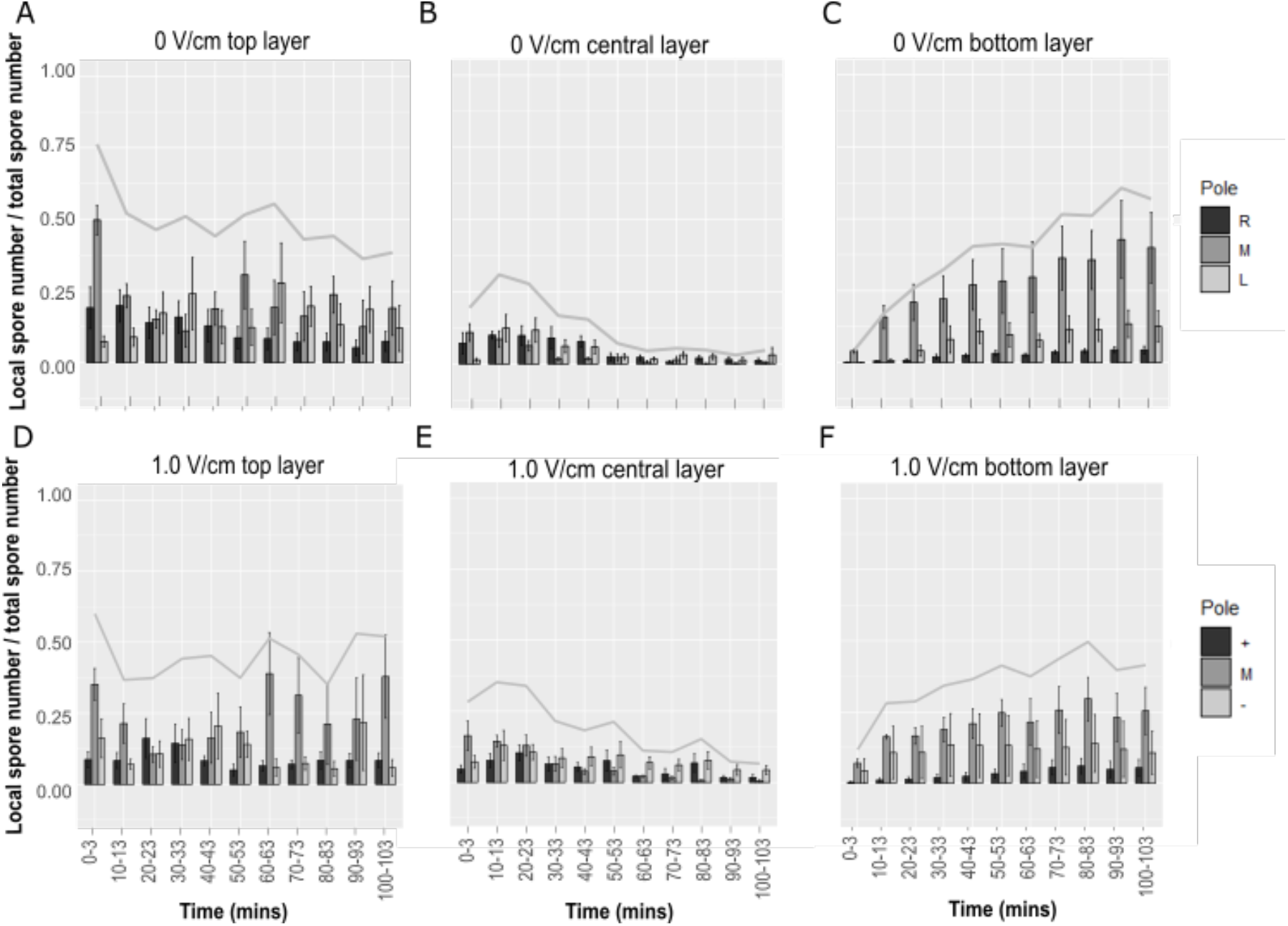
Ionic currents in the medium are necessary for zoospore electrotaxis. Time-course of the distribution of spores detected in the top, central and bottom layers of the medium in the V-slide containing only deionised water. At each timepoint, each bar represents the proportion of spores close to the positive electrode or the right side (dark grey), the negative electrode or the left side (light grey), and in the middle (medium grey); error bars, s.e.p. The grey line is the proportion of all spores detected in that layer, at each timepoint. No significant difference has been observed between the zoospore population at the positive and negative poles, in any of the three layers (p-value between positive and negative > 0.05; t-test, n = 6).

### Exposure to an external electric field does not perturb zoospore encystment

The effects of external electric fields on other phases of the *P. palmivora* life cycle are unknown. Data on fungi, which are morphologically similar, suggest that electric fields could affect spore germination and hyphal growth (McGillivry & Gow, 1986). However, *P. palmivora* zoospore morphology and behaviour are typical of diatoms and have no parallels in fungi (C. E. Bimpong & Hickman, 1975). Therefore, we decided to investigate the effect of external electric fields on all of *P. palmivora*’s early life cycle phases.

Encystment is the process when zoospores lose their flagella, rearrange their organelles, a new cell wall is formed, and adhesive material is secreted to attach to the host (Figure 1 A) (C. E. Bimpong & Hickman, 1975). To test the hypothesis that external fields can accelerate or delay the encystment process, we pipetted *P. palmivora* zoospores near the positive or negative electrode of the V-slide and monitored their vertical position over time. Since encysted spores cannot swim and are known to precipitate at the bottom of liquid media, the ratio between the number of zoospores in the bottom layer and those still in suspension indicates the proportional encystment. We observed no difference in proportional encystment between zoospores exposed to 1.0 V/cm and those not exposed, regardless of their proximity to either electrode (Figure 4 A).

**Figure 4:**
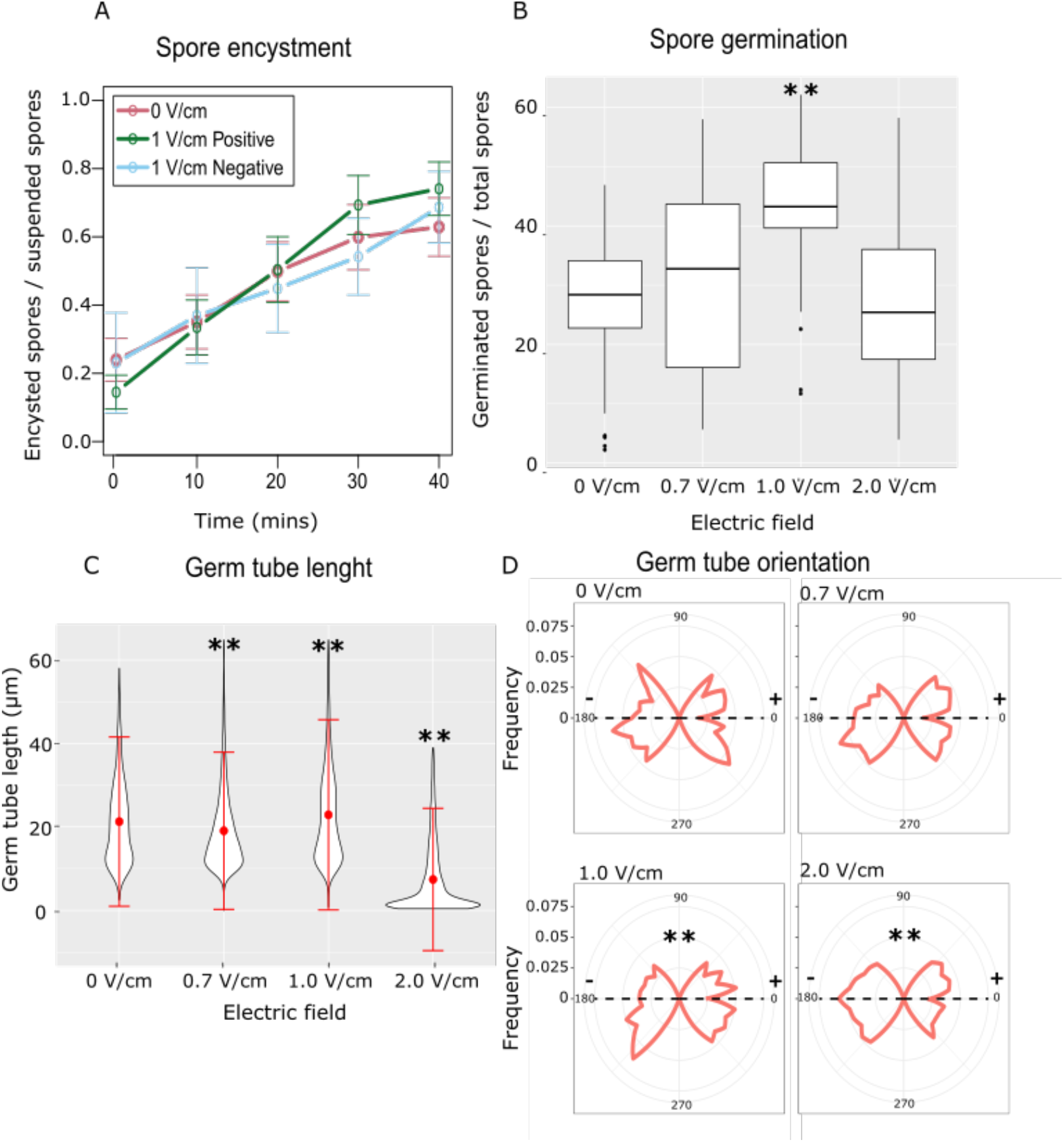
Electric fields do not affect zoospore encystment but affect other aspects of zoospore germination. **A**) Proportional encystment for spores pipetted near the positive (green line) or negative (blue line) electrode; error bars, s.e.m. (n =6). **B**) Distribution of the proportion of spore germination when exposed to various electric fields; ** = p-value < 0.01 between applied field and 0V/cm; Wilcoxon rank sum exact test (n = 3) **C**) Distribution of germ tube length when exposed to various electric fields; ** = p-value < 0.01 between applied field and 0V/cm; Tukey multiple comparisons of means test (n = 3) **D**) Distribution of germ tube orientation observed within ±45° from 0° (*i.e*. towards the positive pole) and within ±45° from 180° (*i.e*. towards the negative pole), when exposed to various electric fields. We compared the two distributions in each electric field: ** p-value < 0.01 between the orientations 0° ± 45° and 180° ± 45°; Wilcoxon rank sum test.

This strongly suggests that the electric field does not affect the biological process of spore encystment.

### Exposure to an external electric field increases zoospore germination rate and alters germ tube growth and orientation

We then asked whether the same electric field could perturb zoospore germination. An encysted spore germinates when small vesicles accumulate at a point at the surface of the cyst. This causes the formation of the germ tube which elongates through the accumulation of more vesicles at the tip (Figure 1 A) (C. E. Bimpong & Hickman, 1975).

First, we used mechanical vibrations to induce encystment (Tokunaga & Bartnicki-Garcia, 1971), a required step before germination. We then exposed the encysted zoospores to 0.7 V/cm, 1.0 V/cm and 2.0 V/cm in the V-slide and imaged them every 10 minutes for 100 minutes. To automate the image processing pipeline, we developed an *ad hoc* ImageJ macro to identify the germinated zoospores and to measure the length and orientation of their germ tube.

Interestingly, we observed a significant increase in spore germination at 1.0 V/cm compared to 0 V/cm (Wilcoxon test, p < 0.01) (Figure 4 B), with a higher proportion of germinated spores close to the negative electrode (Supplementary Figure 5; Wilcoxon test, p < 0.01). We measured a statistically significant decrease in the average germ tube length at 0.7 and 2.0 V/cm and a small but statistically significant increase at 1.0 V/cm, when all compared to 0 V/cm (Figure 4 C; Tukey tests, p < 0.01). Finally, when exposed to 1.0 and 2.0 V/cm, the germ tube orientation was mildly biased towards the negative pole (Figure 4 D; Wilcoxon rank sum test, p < 0.01).

## Discussion

The capability of microorganisms to track plant roots in a complex soil environment rich in chemical and physical stimuli is a poorly understood process. Pathogens and symbionts notably detect and navigate chemical and physical gradients emitted by their targets (Cameron & Carlile, 1978; Steinkellner et al., 2007). In this paper, we characterise the little-studied phenomenon of pathogen electrotaxis and break down how electric cues affect different stages of *P. palmivora*’s life cycle (Figure 1A).

We confirmed that *P. palmivora* zoospores are electrotactic towards the positive pole when exposed to weak electric fields (0.7 – 1.0 V/cm and 31 – 56 μA). Crucially, we showed that in these conditions electrotaxis is dominant over any chemotactic secondary effect and that ionic currents are required. We also showed that the same electric fields do not affect zoospore encystment. Based on these results, we present here a simple model of population dynamics in 3D for zoospores in electric fields (Figure 5). Swimming zoospores tend to accumulate in the top ~ 600 μm of the liquid and move symmetrically in both directions from the centre of the chamber (Khew & Zentmyer, 1974). In presence of the electric field, the zoospores’ movement is biased towards the positive pole, indicating electrotaxis.

**Figure 5:**
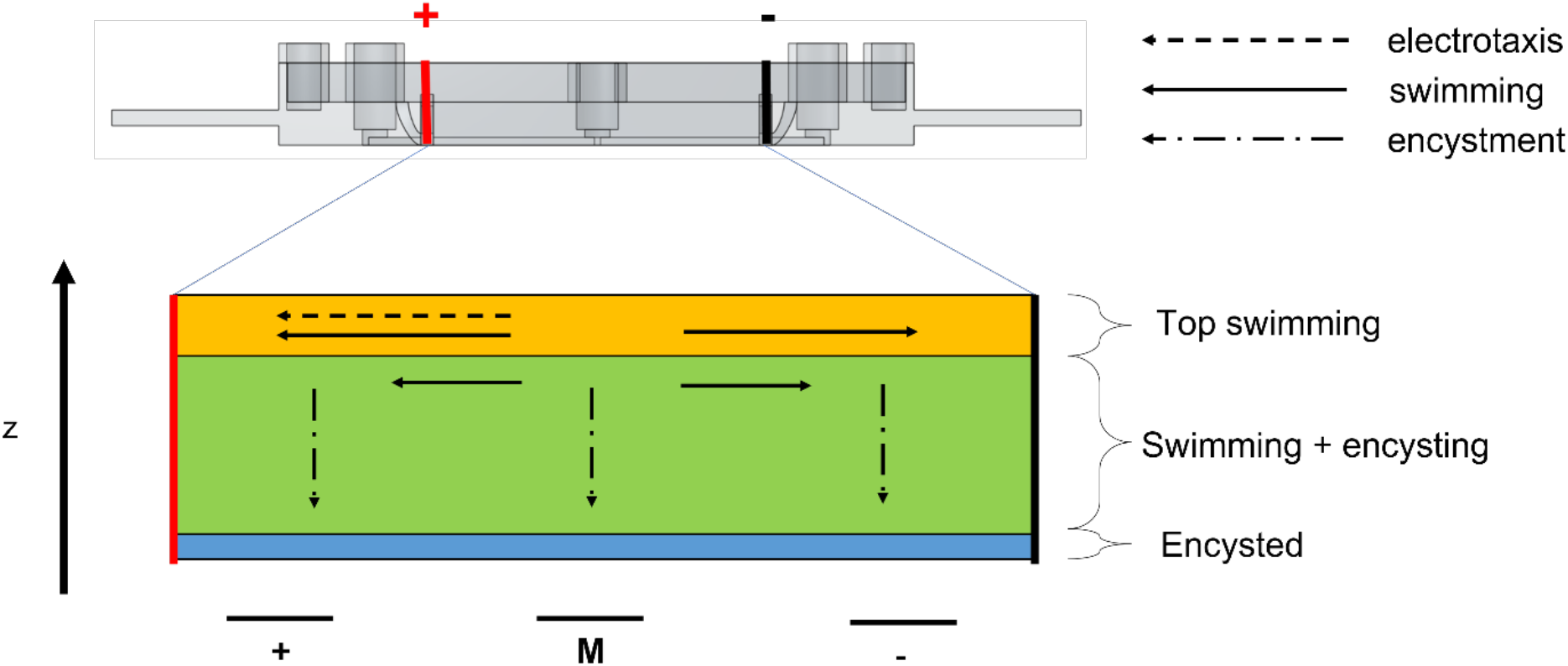
Model of zoospore movement in an electric field *in vitro*. When exposed to an external electric field *in vitro, P. palmivora* swimming zoospores are mostly concentrated in the very top layers of the liquid media. The swimming is biased towards the positive pole (electrotaxis), where the zoospores begin encysting. As zoospores encyst, they precipitate to the bottom of the V-slide.

The mechanism used by the swimming zoospores to detect the electric field remains elusive. Swimming *P. palmivora* zoospores are propelled by the pulling action of a tinsel-like flagellum while a second flagellum is used for steering, resulting in a tumbling motion (Judelson & Blanco, 2005; Morris & Gow, 1993). Since the tinsel-like flagellum is negatively charged and the other flagellum is positively charged (Hardham, 1989; Morris & Gow, 1993), it has been proposed that a simple flagella reorientation due to electrostatic forces could be the main mechanism behind electrotaxis in these spores (Morris & Gow, 1993; Sarkar et al., 2022). If this were the case, an electric field alone with no electrical current or ion flow would be sufficient to trigger the electrotactic behaviour. Instead, here we have shown that when the field is maintained at 1.0 V/cm, but the current is reduced from ~56 μA to ~12 μA (by a significant reduction of ion concentration in the medium), electrotaxis is completely lost. This suggests that ionic currents in the medium are necessary for electrotaxis and that *P. palmivora* zoospores rely on biophysical and molecular mechanisms for sensing such currents.

Although the zoospores maintain the same probability of encystment across the entire chamber even in presence of the electric field, more encysted and encysting spores are found close to the positive pole simply due to the electrotactic behaviour during the early swimming phase.

Since plant roots produce endogenous electric fields and ionic currents of intensities roughly comparable to the ones used in these in vitro experiments (Miller & Gow, 1989; van West et al., 2002), it is plausible that electrotaxis had been selected through evolution to provide *P. palmivora* zoospores with a redundant mechanism to target roots (Toko et al., 1989; van West et al., 2002). However, once their host is reached, it seems that the encystment process is not induced by the same electric fields and currents. Other chemical and physical cues are more likely to trigger this important physiological transition.

Interestingly, we have also shown that, after encystment, an electric field of 1.0 V/cm is sufficient to increase *P. palmivora* zoospore’s probability of germination. This is consistent with the idea that ideally germination should be triggered when the zoospore has encysted on the root’s surface, where its external electric field and ionic currents are the strongest.

We observed that, when exposed to field intensities higher than 1.0 V/cm, the germ tubes preferentially grow towards the negative pole, indicating some degree of electrotropism. We speculate that 1.0 V/cm might be close to the minimum intensity required to redirect the vesicles depositing membrane at the hyphal tip (C. E. Bimpong & Hickman, 1975), similar to what shown in pollen tube growth and *Neurospora crassa* hyphae (Malhó et al., 1992; McGillivray & Gow, 1986). However, the decrease in germ tube length and lack of increased germination at 2 V/cm, where the effect is stronger, might also indicate a stress response. Therefore, a more detailed characterization seems necessary to fully understand germ tube electrotropism in *P. palmivora*’s zoospores.

Overall, we characterized *P. palmivora* zoospore electrotaxis in detail and described the effects of electric fields on cyst germination *in vitro*. Future work should focus on investigating these phenomena *in vivo* and investigate the zoospores’ electric field and current sensing mechanisms.

## Materials and Methods

### *P. palmivora* and culture conditions

*P. palmivora* YFP (P16830 LILI YFP KDEL) mycelia were cultured on V8 medium (100 ml/l V8 juice, 1 g/l CaCO2, 50 mg/l β-sistosterol, 15 g/l bacteriological agar, 50 μg/ml G418) at 28°C in constant light. The zoospores were released and collected by flooding a 6-day-old plate with 5 mL of 1/500X liquid MS medium (1X MS medium is 5 g/l Sucrose, 8.6 mg/l MS salts, 0.5 g/l MES buffer, adjusted to pH 5.7 with 1M Tris HCL) and incubating for one hour at 22°C.

### Design and fabrication of the V-slide

The V-slide was designed using the CAD software OnShape and printed with PLA filament using an Ultimaker 2+ 3D printer. A coverslip was adhered to the base of the V-slide with silicon and left to dry overnight. Two platinum-iridium foils (3.9 mm × 6 mm) were soldered along the long edge to a copper wire which was fed through the two 1 mm holes above the electrotaxis chamber. This allows the plate electrodes to be held in place and connected to a PS-1302 D power supply (Voltcraft, UK) without any contact between the media and any non-platinum iridium circuit components (Figure 2 and Supplementary Figure 1).

### Electrotaxis experiments

The electrotaxis experiments were performed by mounting the V-slide on a Nikon Eclipse-Ti inverted microscope. The electrodes were coated with 1% Agarose in 1/500X MS medium, 1 mL of 1/500X MS medium was added to the chamber and 50 μl of spore solution (20,000 spores/ml) was pipetted in the middle of the chamber. The electric field was then turned on generating fields (and corresponding measured currents) of 0.5 V/cm (24 ± 10 μA), 0.7 V/cm (31 ± 8 μA) or 1.0 V/cm (55 ± 14 μA). Stacks (step size 30 μm) were taken at 10 min intervals for 100 min in three regions: next to each electrode and in the middle of the V-slide (Figure 2A). Swimming and encysted spore numbers were then counted using an existing automated ImageJ macro (Schneider et al., 2012). The counted spores were then manually divided into three layers: the top layer being the first 540μm of the liquid media, or the top 18 slices in the stack (Supplementary Figure 2), the bottom as the single slice where encysted spores are in focus and the central layer as the all the space in-between. The proportion *p*_*r,l*_ of spores in each position (3 regions x 3 layers) was calculated as:

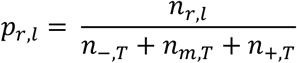

Where: *n_r,l_* is the number of spores at position (*r*, *l*); *r* = {−, *m*, +} for the negative, middle and positive regions; *l* = {*t*, *c*, *b*} for the top, central and bottom layers; *n_r,T_* = *n_r,t,_* + *n_r,c,_* + *n_r,b_*

### Chemotaxis experiments

The chemotaxis experiments were performed by adding 8 g/l of sucrose to the agarose coating of only one of the electrodes. The experiment was run in the same manner as the electrotaxis experiments but without connecting the electrodes to the power supply. The zoospores were counted, and their distribution was calculated in the same manner as for the electrotaxis experiments.

### Pre-exposure test

The pre-exposure experiments were performed by exposing the media in the V-slide to 1 V/cm for 100 minutes. The field was then switched off, the zoospores were added, and imaged as during the electrotaxis experiments. The zoospores were counted, and their distribution was calculated in the same manner as for the electrotaxis experiments.

### Encystment experiments

The encystment experiments were performed similarly to the electrotaxis experiments, but with the spores pipetted near one of the poles. The proportion *pe_r_* of spores encysted at each pole was calculated as:

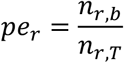

Where *n_r,b_* is the number of zoospores laying at the bottom of the slide, at the poles *r* = {−, +}; *n_r,T_* = *n_r,t,_* + *n_r,c,_* + *n_r,b_* with {*t*, *c*, *b*} for the top, central and bottom layers.

### Germination experiments

Once collected as described above, the zoospores were vortexed for 30 seconds and incubated at room temperature for 4 minutes to cause encystment without inhibiting germination (Tokunaga & Bartnicki-Garcia, 1971). The zoospores were then pipetted to obtain an even distribution across the V-slide and exposed to the electric field for 100 minutes. Images of the bottom layer of each region were taken on the Nikon Eclipse-Ti inverted microscope every 10 minutes. The number of germinated spores, length of the germ tube and direction of the germ tube were quantified using an ImageJ macro developed for this project and available on request. The proportion of germinated spores over total encysted spore numbers was then calculated.

### Statistical analysis

When comparing two samples of measurements, each distribution was first checked for normality with the Shapiro–Wilk normality test with alpha-level = 0.05. Normal distributions were tested with the two-tails Student’s t-test without assuming equal variances (Welch two-sample t-test).

When comparing multiple samples of measurements, we used Shapiro–Wilk normality test to check for normality with alpha-level = 0.05. Normal distributions were tested with one-way ANOVA followed by Tukey multiple comparisons of means test; if one of the two distributions was not normal, the Kruskal-Wallis rank sum test was used followed by the Wilcoxon rank sum exact test. All statistical tests were performed with R.

## Supporting information

Supplemental Informations

## Acknowledgements

We thank S. Schornack for providing *P. palmivora* and the invaluable support during the project. We also thank M. Barkoulas and the FILM facility at Imperial College London for the use of their microscopes.

## Notes

### Competing Interest Statement

The authors have declared no competing interest.

