## Supplemental Informations for "Enhanced germination and electrotactic behaviour of *Phytophthora palmivora* zoospores in weak electric fields"

### Supplementary Information

| | FP (%) | FN (%) |
|---|---|---|
| **Encysted spore macro** | 7.73 ± 0.28 | 3.74 ± 0.07 |
| **Swimming spore macro** | 1.29 ± 0.03 | 1.86 ± 0.03 |

**Supplementary Table 1: Validation of macros performance.** Percentage of false positives (FP = spores identified by the macro, but not confirmed by a human) and false negatives (FN = spores identified by human but missed by the macro); total number of images, n = 60

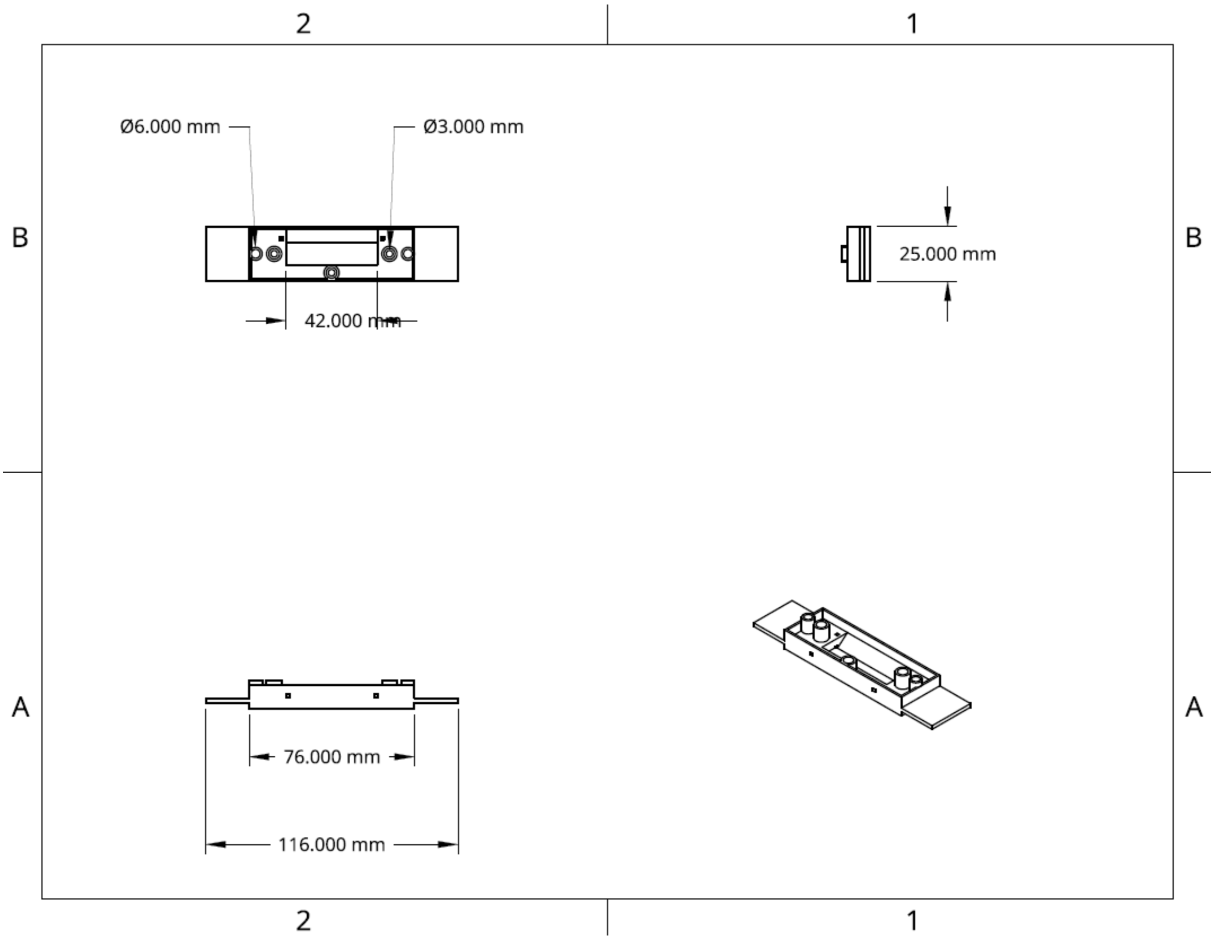


**Supplementary Figure 1: V-slide design.** Summary of the CAD schematic used to 3D print the V-slide.

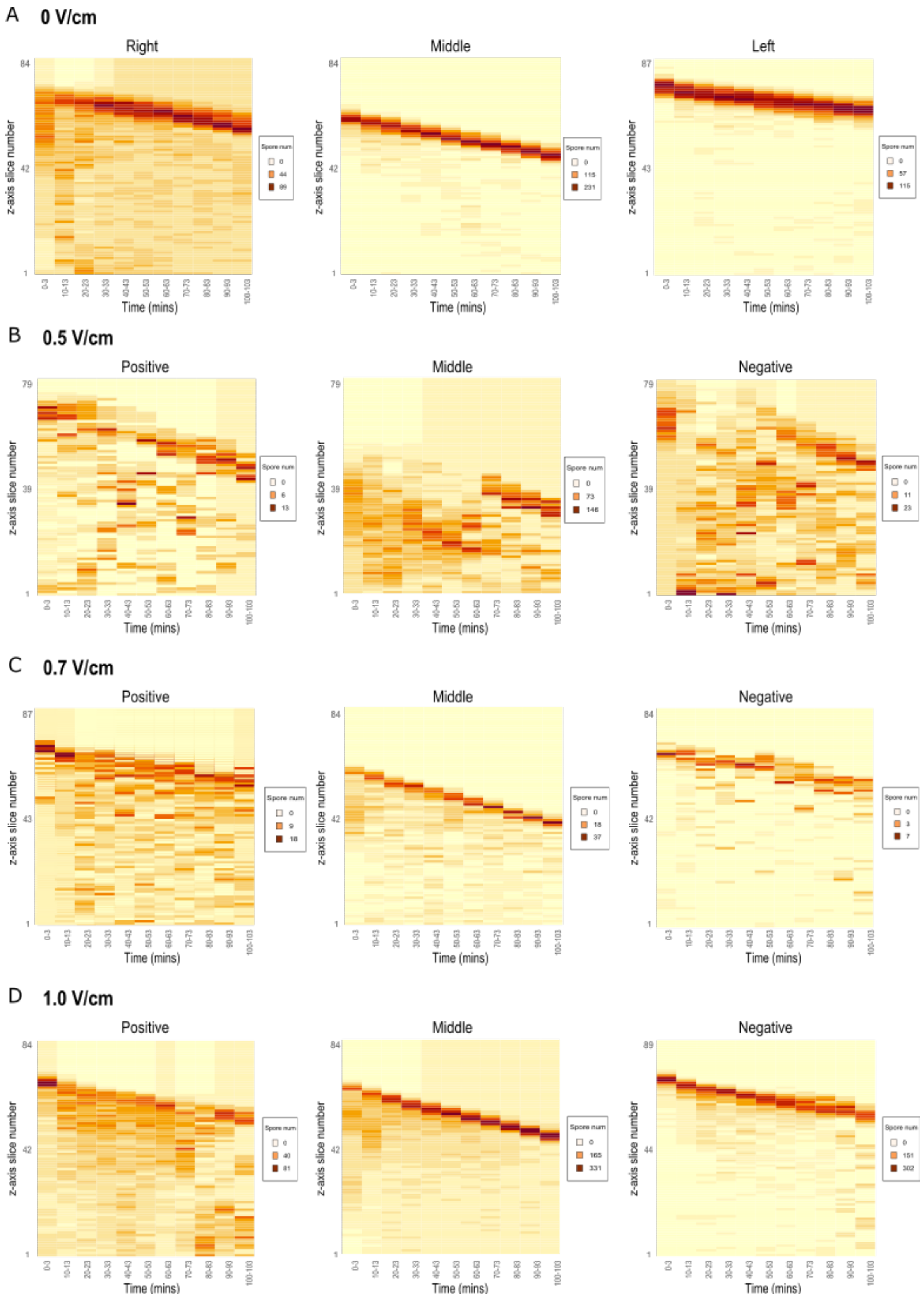


**Supplementary Figure 2: Vertical zoospore distribution in the V-slide.** Each panel is a representative heatmap showing the number of spores counted in each 30µm-thick z-slice (Y-axis), at a given time-point (X-axis), from a single repeat. Spore counting was limited to three regions of the V-slide: close to the positive pole (Positive), in the middle of the slide (Middle) and close to the negative pole (Negative). Data is shown for 0 V/cm (**A**), 0.5 V/cm (**B**), 0.7 V/cm (**C**) and 1.0 V/cm (**D**) electric fields.

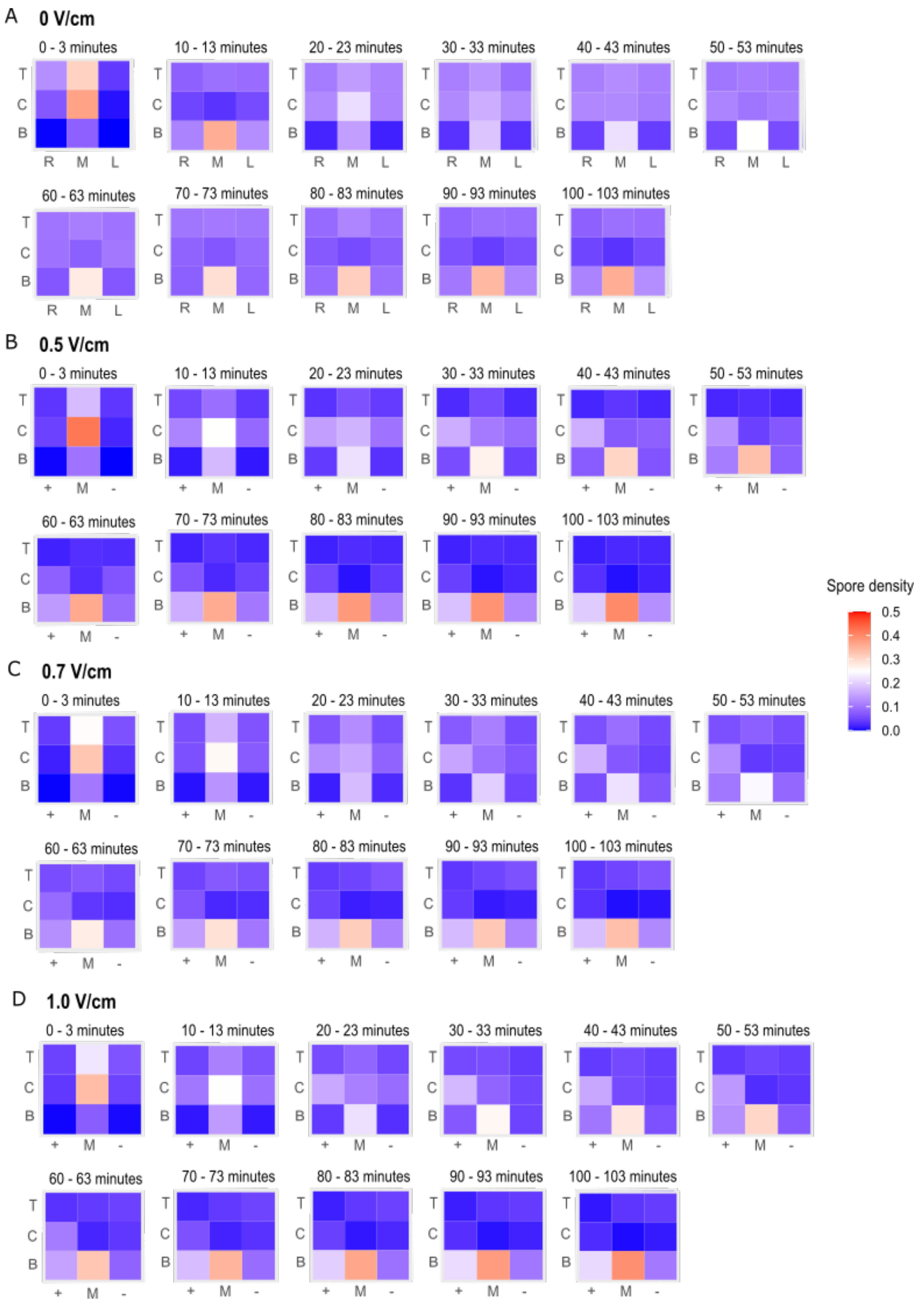


**Supplementary Figure 3: 3D zoospore distribution in the V-slide.** Each panel is a heatmap showing the mean proportion of zoospores detected in the top (T), central (C) and bottom (B) layers, and close to the positive (+) or negative (-) poles and in the middle of the V-slide (M). Each heatmap shows the 3D zoospore distribution in the given time interval. Each zoospore proportion is calculated as the number of spores detected in the sub-volume divided by the total number of spores detected in the V-slide in the same time interval. Data is shown for 0 V/cm (**A**), 0.5 V/cm (**B**), 0.7 V/cm (**C**) and 1.0 V/cm (**D**) electric fields (n = 9).

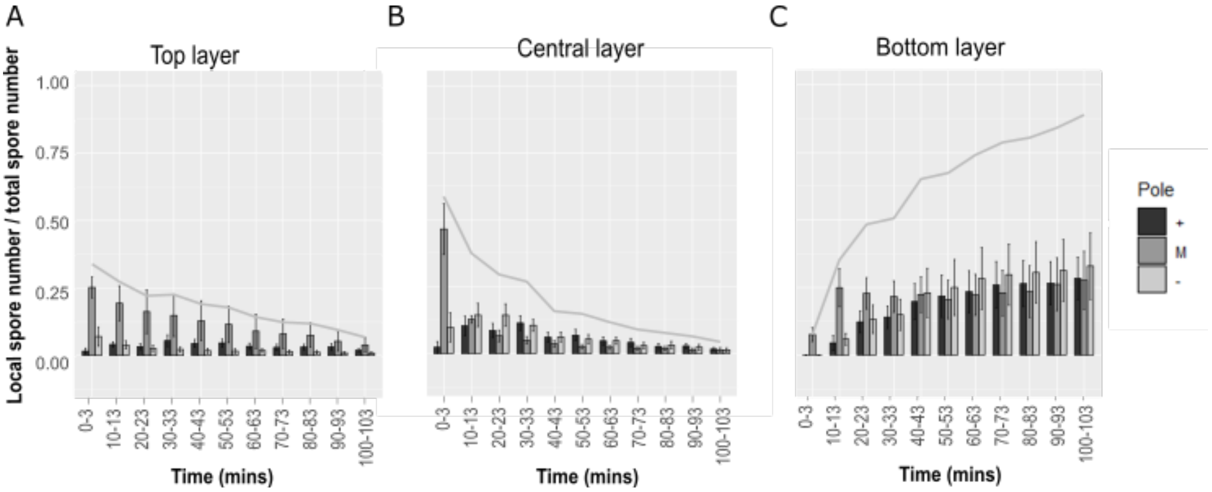


**Supplementary figure 4: Zoospore distribution in pre-exposed medium.** Distribution of spores detected in the top (A), central (B) and bottom (C) layers of the medium in the V-slide. Bars represent the proportion of spores in each region of the V-slide: close to the positive electrode or right side (dark grey), the negative electrode or left side (light grey), and in the middle (medium grey); error bars, s.e.p. The grey line is the proportion of all spores detected in that layer, at each timepoint. Only a symmetric movement is observed, without any significant bias of the distribution toward any of the poles (n=6).

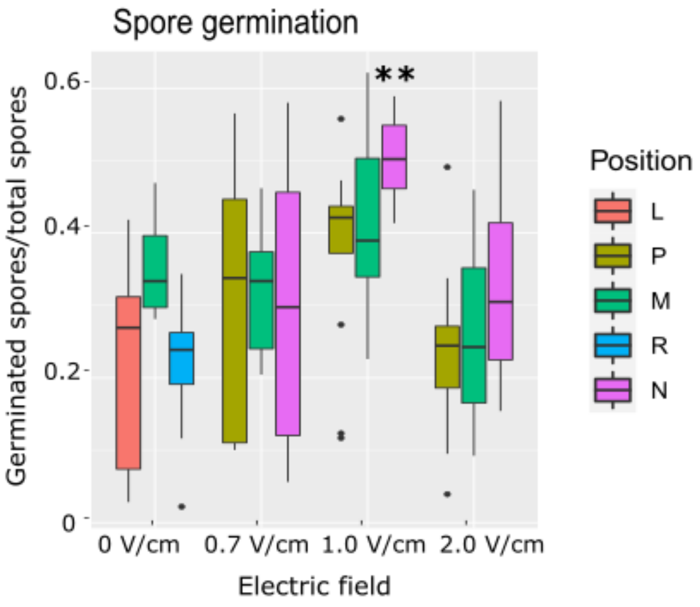


**Supplementary Figure 5: Effects of electric fields on spore germination and germ tube growth. A**) The percentage of spore germination increases at 1.0 V/cm close to the negative pole. The V-slide regions are labelled positive (P) and negative (N) when the field is on, left (L) and Right (R) when the field is off, and middle (M) in both cases (** = p-value < 0.01 between positive and negative; Wilcoxon test, n = 3).
